# A genetically encoded fluorescent sensor for *in vivo* imaging of GABA

**DOI:** 10.1101/322578

**Authors:** Jonathan S. Marvin, Yoshiteru Shimoda, Vincent Malgoire, Marco Leite, Takashi Kawashima, Thomas P. Jensen, Erika L. Knott, Ondrej Novak, Kaspar Podgorski, Nancy J. Leidenheimer, Dmitri A. Rusakov, Misha B. Ahrens, Dimitri M. Kullmann, Loren L. Looger

## Abstract

Current techniques for monitoring GABA, the primary inhibitory neurotransmitter in vertebrates, cannot follow ephemeral transients in intact neural circuits. We applied the design principles used to create iGluSnFR, a fluorescent reporter of synaptic glutamate, to develop a GABA sensor using a protein derived from a previously unsequenced *Pseudomonas fluorescens* strain. Structure-guided mutagenesis and library screening led to a usable iGABASnFR (ΔF/F_max_ ~ 2.5, K_d_ ~ 9 μM, good specificity, adequate kinetics). iGABASnFR is genetically encoded, detects single action potential-evoked GABA release events in culture, and produces readily detectable fluorescence increases *in vivo* in mice and zebrafish. iGABASnFR enabled tracking of: (1) mitochondrial GABA content and its modulation by an anticonvulsant; (2) swimming-evoked GABAergic transmission in zebrafish cerebellum; (3) GABA release events during inter-ictal spikes and seizures in awake mice; and (4) GABAergic tone decreases during isoflurane anesthesia. iGABASnFR will permit high spatiotemporal resolution of GABA signaling in intact preparations.

## Introduction

γ-Amino butyric acid (GABA) is a ubiquitous inhibitory neurotransmitter (the principal one in vertebrates), reducing neuronal excitability by receptor-mediated hyperpolarization of membrane potential and shunting of excitatory currents. GABA can also be excitatory in early development and in cell types with high intracellular chloride^1^. GABA_A_ receptors are hetero-pentameric Cl^-^-conducting channels^2^. GABA_B_ receptors are hetero-dimeric metabotropic receptors that are coupled to multiple ionic currents by G_i/o_-proteins^3,4^, and are more sensitive to GABA than most synaptic GABA_A_ receptors^5^. The inhibitory role of GABA and its receptors has been the subject of significant pharmacological intervention. Drug classes include GABA_A_ receptor allosteric modulators (*e.g.* sedative, anxiolytic and anticonvulsant benzodiazepines), GABA_B_ agonists (*e.g.* the anti-spastic baclofen), GABA reuptake inhibitors (*e.g.* the anticonvulsant tiagabine), and GABA transaminase inhibitors (*e.g*. the antiepileptic vigabatrin). GABA receptor antagonists include both drugs and toxins.

Despite the clinical efficacy of drugs that augment GABAergic neurotransmission, the details of GABA release from neurons, diffusion to both post-and pre-synaptic receptors, reuptake by neurons and astrocytes, recycling through the glutamate/GABA-glutamine shuttle^6^, and other critical mechanisms, including its role in metabolism, remain largely unknown. GABA signaling has traditionally been inferred from a combination of electrophysiology and pharmacology^7^. Direct detection of GABA has usually been achieved by microdialysis followed by derivatization and quantification by HPLC/electrochemical detection^8^. In addition to the inherently poor spatiotemporal resolution (on the order of minutes and hundreds of microns) and the sensitivity loss from derivatization, direct detection of GABA is further limited by its being about 20-fold less abundant than glutamate^9^.

While genetically encoded fluorescent sensors for calcium^10,11^ have been optimized for imaging neural activity, and sensors for the excitatory neurotransmitter glutamate^12^ have been developed, methods for directly imaging GABAergic signaling are lacking. Recently, a semi-synthetic fluorescent sensor for GABA was reported^13^, based on the attachment of organic fluorophores to the GABA_B_ receptor. However, its utility in neuronal systems is limited, since it requires the addition of two separate small molecules and its affinity for GABA is extremely weak (~400 μM). Furthermore, over-expression of GABA receptors could disrupt multiple aspects of cellular signaling, and similarly, hetero-oligomerization with endogenous receptor pools could degrade sensor functionality. Alternatively, inhibitory neurotransmission can be indirectly inferred from intracellular chloride concentration [Cl^−^]_i_, measurable with Clomeleon and improved variants^14^. However, chloride sensors have poor biophysical performance; [Cl^−^]_i_ does not change much during inhibition; the sensors conflate the activities of all ligand-gated (GABA, glycine) and voltage-gated Cl^−^ channels; and GABA_B_ receptors are G_i/o_-coupled receptors that do not directly affect [Cl^−^]_i_. Thus, direct detection is preferable for inference of GABAergic signaling in general, and is required for mechanistic studies of synaptic input-output transformations, as has been done recently with glutamate release and presynaptic Ca^2+^ imaging^15^.

We previously reported a method for developing genetically encoded fluorescent sensors with excellent ligand-binding specificity, affinity, and kinetics using bacterial periplasmic binding proteins (PBPs)^16^. This approach to sensor development couples the Venus flytrap-like conformational change that occurs upon ligand binding to changes in the fluorescence of an inserted circularly permuted fluorescent protein. We previously used this technique to develop a glutamate sensor^12^ for *in vivo* imaging. Here we report similar development of an intensity-based GABA Sensing Fluorescence Reporter (iGABASnFR) and illustrate its utility for observing GABA uptake into mitochondria, and GABA release in cultured neurons and acute mouse brain slice. *In vivo*, we observed bulk GABA transients in mouse visual cortex, and GABA release synchronized with inter-ictal spikes in a mouse model of epilepsy. Finally, in zebrafish we used iGABASnFR to correlate GABAergic signals across the cerebellum to motor output.

## Results

### Sensor engineering

There are many potential scaffolds for developing a GABA sensor. We deemed the GABA_A_ and GABA_B_ receptors inappropriate since the GABA-binding sites of the former are at protein-protein interfaces^2^, without an obvious route for coupling GABA-dependent conformational changes to effects on a fused fluorescent protein, and the latter is difficult to express heterologously^17^. Furthermore, as discussed above, over-expression of these receptors could alter cell and sensor function in many ways. In theory, one could redesign the binding site of an existing sensor (*e.g.* iGluSnFR) to make it specific for GABA. Our attempts to do so failed (data not shown).

GABA is a prominent signaling molecule in the plant-soil interface; in fact, it was first isolated from potato tubers^18^. *Agrobacterium tumefaciens* is a plant pathogen that expresses two periplasmic GABA-binding proteins. Atu2422 binds GABA and other small amino acids^19^. While we were able to develop a sensor from this scaffold, it has very weak (high-μM) affinity for GABA, and orders-of-magnitude tighter affinity (low-μM) for alanine and glycine (**Supp. Fig. S1**). Its weak, nonspecific response to GABA makes it an inappropriate sensor scaffold. Atu4243 has more stringent specificity and tighter affinity for GABA (9 μM)^20^. Unfortunately, Atu4243 with either cpGFP or cp-superfolderGFP (cpSFGFP) inserted – at locations that we planned to optimize into sensors – did not translocate to the membrane surface in cultured cells when cloned into the pDisplay vector (Invitrogen) (**Supp. Fig. S2**) and was abandoned as a scaffold.

Prior to the structures of Atu2422 and Atu4243 being published, a strain of *Pseudomonas fluorescens* (ATCC #BAA1781) was reported to rapidly uptake GABA^21^, presumably through a high-affinity PBP/transporter. We received a culture of this strain (CNG89; a gift from Dr. Catherine Guthrie) and sequenced its genome. We identified “Pf622” as a homologue of Atu4243 (**Supp. Fig. S3**) and cloned it by PCR into the pRSET vector (Invitrogen) for heterologous expression in *Escherichia coli*. Its affinity was not in the low-nM range as expected from the original publication^21^, but rather 130 μM as determined by isothermal titration calorimetry (**Supp. Fig. S4a**). (The calculated molecular weight of Pf622 is 37 kDa, and 35 kDa after cleavage of the predicted periplasmic leader peptide. This is smaller than the apparent 42 kDa determined by SDS-PAGE for the presumed high-affinity GABA-PBP isolated from CNG89^22^.) A fluorescent allosteric signal transducer^23^, introduced by coupling an environmentally sensitive fluorophore, JF-585^24^, *via* maleimide-thiol chemistry to a cysteine mutation at a residue in the hinge of the protein (V278C), resulted in a hybrid sensor with 240 μM affinity for GABA (**Supp. Fig. S5**). Stopped-flow kinetic analysis revealed atypical kinetics. Instead of a single exponential rise in fluorescence upon mixing of protein and GABA, the data is best fit by a double exponential (**Supp. Fig. S6a,b**), suggesting multiple steps link GABA binding to fluorescence change. Regardless, the rise time for Pf622.V278C-JF585 binding GABA is longer than it is for iGluSnFR binding glutamate (**Supp. Fig. S6c**).

Pf622 with cpGFP or cpSFGFP inserted at any of several sites translocated to the membrane when cloned into pDisplay and expressed in mammalian cells (**Supp. Fig. S7**). The improved expression and photostability observed by the substitution of cpSFGFP for cpGFP in iGluSnFR^25^ led us to use the former in this work. Based on its homology to Atu4243, we expected insertion of cpSFGFP after residue 276 of Pf622 (sequence info in **Supp. Fig. S8**) to yield a good sensor after optimization of the residues linking the binding protein and the FP. Linker optimization yielded a version of the sensor (L1-LA/L2-AN) for which we solved a crystal structure in the unliganded/open state (**Supp. Fig. S9**). In conjunction with its homology to Atu4243, for which both the unliganded/open and bound/closed structures are available^20^, we tested mutations to the hinge region of Pf622 that we expected to allosterically modulate affinity^26^. Hinge mutation F101L greatly tightened affinity (**Supp. Fig. S10**). Mutation of a residue in Pf622 (N260A) that potentially interacted with cpSFGFP, and was also a potential glycosylation site, substantially increased brightness in HEK cells (**Supp. Fig. S11**). Re-optimization of the linkers (L1-LAQVR, L2-AN) plus a mutation in the circularly permutated strand of SFGFP (F145W), resulted in a variant, called iGABASnFR, worthy of further characterization. iGABASnFR is Pf622 with cpSFGFP inserted after residue 276, with optimized linkers (L1-LAQVR, L2-AN), a hinge mutation (F101L), an interface/expression mutation (N260A), and an additional GFP mutation (F145W).

### In vitro *characterization*

Purified iGABASnFR shows a maximum ΔF/F of ~ 2.5 and a *K*d for GABA of ~ 9 μM (**Supp. Fig. S12a**). It shows no affinity for other amino acids except very weak affinity for glycine (500 μM), alanine (830 μM), and histidine (2.4 mM) (**Supp. Fig. 12b**). It has no affinity for similar four-carbon metabolites fumarate, malate, oxaloacetate, nor succinate (data not shown). While we characterized iGABASnFR in various cellular systems, we made additional efforts to increase its affinity and ΔF/F. Mutation of a GABA-binding pocket residue to its Atu4243 homologue, F102Y, had minimal effect on GABA affinity and decreased ΔF/F. Meanwhile, mutation Y137L increased ΔF/F, at the cost of poor expression. The double mutant F102Y.Y137L expressed well, exhibited higher ΔF/F (3.5 vs. 2.5 for iGABASnFR) but weaker affinity (70 μM vs. 9 μM for iGABASnFR; **Supp. Fig. S12a**). Addition of F102G to iGABASnFR (without changes to Y137) also showed higher ΔF/F (4.5) and affinity of 50 μM. These three variants were further characterized.

We first profiled the variants for binding to GABAergic drugs (**Supp. Fig. S12c-f**). Baclofen (β-(4-chlorophenyl)-GABA) affected all three iGABASnFR variants tested (as well as the negative control cpSFGFP, **Supp. Fig. S12c**), whereas vigabatrin (γ-vinyl-GABA, Sabril®) bound to only the F102G variant. Importantly, most drugs showed much weaker binding to the sensors than GABA, allowing them to be used in conjunction with iGABASnFR imaging.

### GABA processing in mitochondria

After being transported into mitochondria (by an as yet unidentified transporter^27^), GABA is degraded by GABA transaminase (GABA-T). Inhibition of GABA-T elevates synaptic levels of GABA^28^ and, of clinical significance, the GABA-T inhibitor vigabatrin is a currently prescribed antiepileptic^29^. We transfected cultured prostate cancer cells (LNCaP) with iGABASnFR.F102Y.Y137L (which doesn’t bind vigabatrin; **Supp. Fig. S12f**), fused to an N-terminal mitochondrial matrix targeting motif from Cox8a (mito-iGABASnFR.F102Y.Y137L). Basal fluorescence in these cells was very low (**Supp. Fig. S13**), consistent with the observation that these cells contain little GABA^30^. In the presence of exogenously applied GABA, robust fluorescence was observed within the mitochondria as expected (**Fig. 1a**). Treatment with varying amounts of GABA increased fluorescence (**Fig. 1b, Supp. Fig. S13**), reaching equilibrium after 48 hours, a time-course presumably reflecting both increasing mito-iGABASnFR expression and continued GABA uptake. The [GABA] dependence of mitochondria-localized iGABASnFR is approximately linear (**Fig. 1c**). Vigabatrin treatment tends to increase iGABASnFR fluorescence in the mitochondria (**Supp. Fig. S14**). To promote endogenous GABA synthesis, LNCaP cells were transfected with a plasmid encoding glutamic acid decarboxylase 1 (GAD1, *a.k.a.* GAD67). GAD1 expression increased the fluorescence of mito-iGABASnFR.F102Y.Y137L within the first and subsequent time-points (50 hours post-transfection) (**Fig. 1d**).

**Fig. 1.**
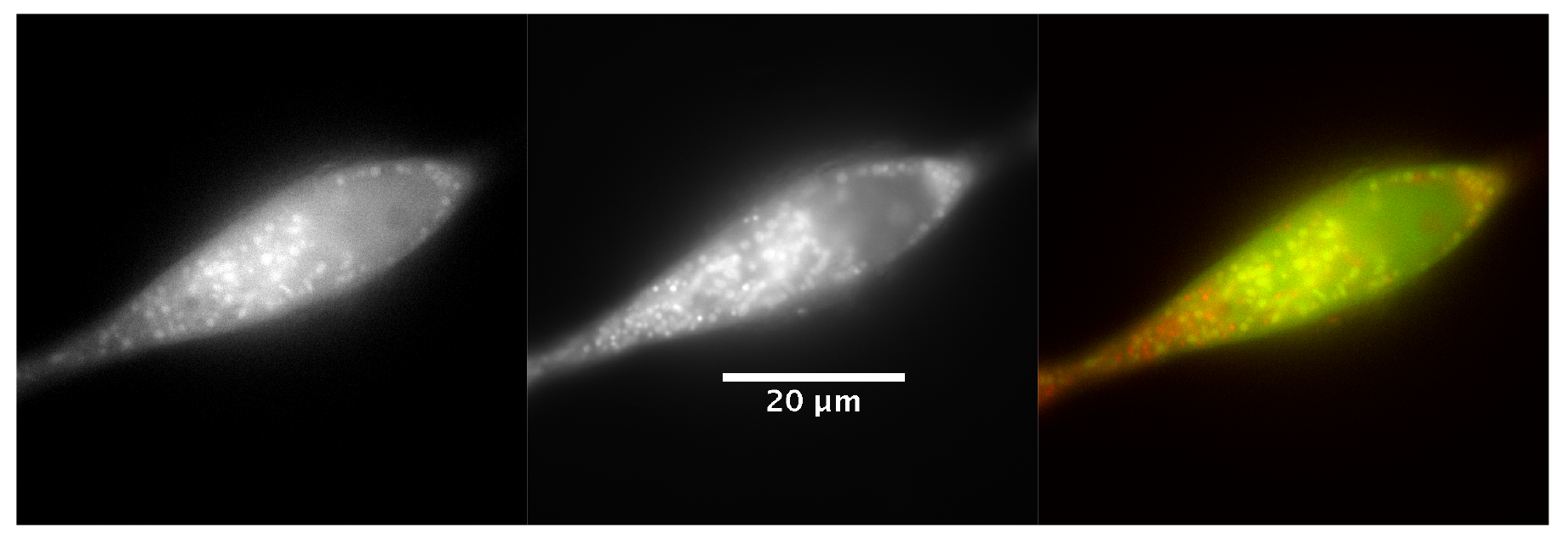
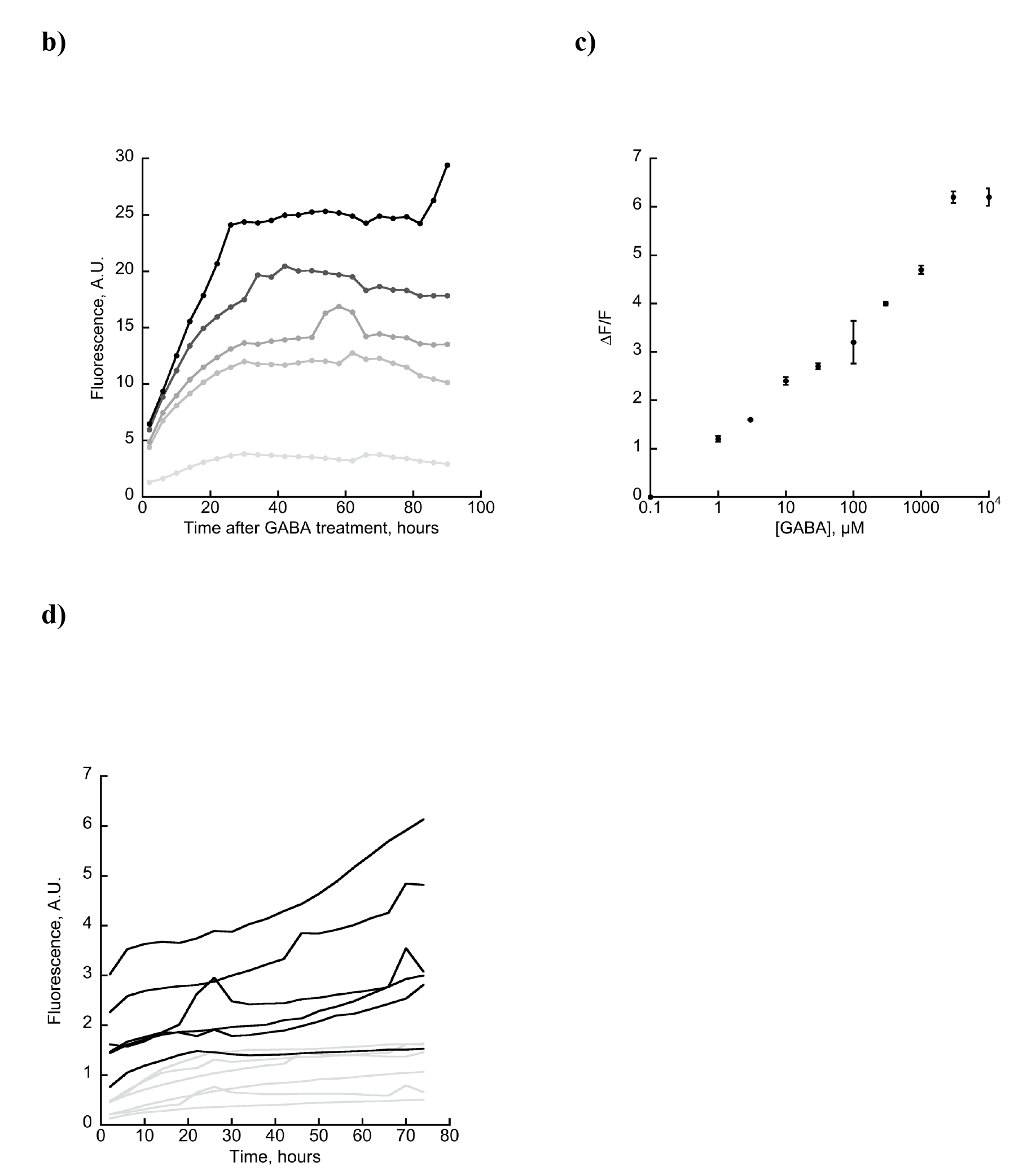
Transport of GABA into the mitochondria is detected by mito-iGABASnFR.F102Y.Y137L. a) LNCaP cells were transfected with mito-iGABASnFR.F102Y.Y137L in the presence of 100 μM GABA. 24 hours after transfection, cells were incubated with MitoTracker Red. Epifluorescence (60x objective) shows co-localization of the two signals. *Left*, green channel. *Middle*, red channel. *Right*, merged. b) LNCaP cells were transfected with mito-iGABASnFR and the media supplemented with GABA 24 hours post-transfection. Fluorescence was measured 2 hours after treatment, and every 4 hours thereafter. b) Time course of fluorescence for different GABA treatments (0, 1, 10, 100, 1000, and 10,000 μM GABA, light grey to black, respectively). c) [GABA] dependence of response was determined by averaging three time points (46, 50, 54 hours) within a single experiment. d) Fluorescence of LNCaP cells transfected with mito-iGABASnFR.F102Y.Y137L and a vector expressing glutamate decarboxylase 1 (GAD1, *a.k.a.* GAD67) relative to an empty vector control (*n*=6). Fluorescence was measured beginning 50 hours post-transfection. Black, GAD1; grey, empty vector. For all plots, fluorescence was quantified with thresholding using an automated algorithm (see **Methods**).

### Neuronal culture characterization

HEK cells transfected with iGABASnFR variants cloned into a modified version of pDisplay lacking the HA tag (pMinDis)^12^ showed good membrane localization (**Supp. Fig. S11**). Primary culture of hippocampal neurons from P0 newborn rats infected with AAV2/1. *hSynapsin1*.iGABASnFR showed good membrane localization and brightness at 14 DIV, but the +F102G and +F102Y.Y137L variants were noticeably dimmer, and showed accumulation in the endoplasmic reticulum (**Supp. Fig. S15**).

Electrical field stimulation (50 Hz) of the cultured neurons produced fluorescence changes from single stimulus-evoked action potentials and increased until plateauing after 40 AP (**Fig. 2a-c**). The response was similar to that of SF-iGluSnFR.A184V^25^ (**Fig. 2d**), except that the magnitude of response was ten-fold lower and more localized (**Supp. Fig S16**), consistent with GABAergic neurons representing a smaller fraction (~10%) of the total neuronal population in hippocampal culture^31^

**Fig. 2.**
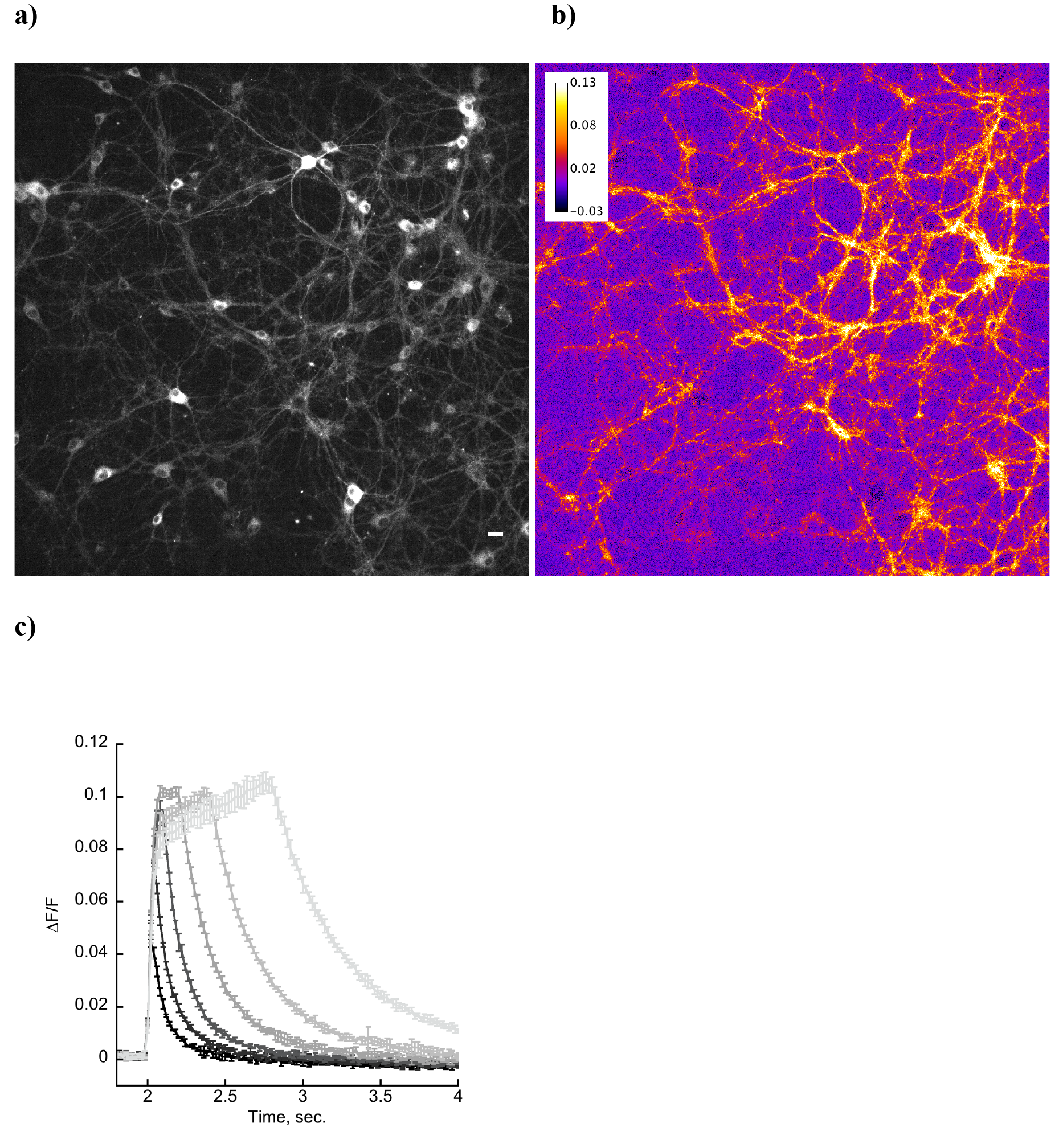
Response of iGABASnFR.F102G in rat hippocampal culture 14 DIV to whole field electrical stimulation. a) Fluorescence image. b) Heat map showing ΔF/F in response to 10 AP stimulation. c) Averaged response (mean ± std.dev., *n*=3) of an ROI selected in a region of maximal ΔF/F to multiple stimuli (50 Hz). Black to light grey: 1, 2, 5, 10, 20, 40 field stimuli. SF-iGluSnFR.A184V responses performed in side-by-side cultures are in **Supp. Fig. S16**. Scale bar 20 μm.

### Hippocampal acute hippocampal slice

To confirm that iGABASnFR remains functional in organized brain tissue, we tested iGABASnFR in hippocampal acute slices prepared from mice with iGABASnFR expressed primarily in pyramidal cells through *in vivo* AAV injection into the hippocampal CA1 region (see Supp. Methods). Although it was challenging to reliably record GABA release with iGABASnFR at the level of individual activated synapses (not shown), as reported previously with iGluSnFR^15^, the sensor displayed excellent S/N ratio in response to paired pulse extracellular field stimulation of the Schaffer Collateral pathway, a protocol used widely in synaptic physiology experiments. iGABASnFR responses to extracellular GABA transients could be recorded from apical dendritic segments as short as 10–15μm traced from the CA1 pyramidal cell body (**Supp. Fig. S17**). Increasing the concentration of extracellular calcium increased the amplitude of fluorescence, consistent with changes in pre-synaptic release probability^32^. It is important to note the value of a spatially resolvable readout of synaptic GABA release given the highly organized and cell type specific nature of GABAergic innervation onto pyramidal cells^33^. Imaging of GABA transients in different cellular compartments enables a high-throughput method to study GABA release from different populations of hippocampal interneurons. Combined with patch-clamp electrophysiology and Ca^2+^ imaging with red-shifted indicators, iGABASnFR could provide a tool for directly studying the local signaling requirements underlying important physiological phenomena such as depolarization induced suppression of inhibition^34^.

### Volume detection of GABA in mouse visual cortex

Tonic GABA release has recently become appreciated as a mechanism for fine-tuning of circuit function^35^, such as supporting context-dependent behavior in a morphologically fixed cortical circuit^36^. In the visual cortex, GABAergic inhibition^37^ and disinhibition^38^ contribute to context-dependent processing and can enhance the distinctness of sensory responses. Hypnotic effects of volatile anesthetics have been associated with GABAergic transmission^39^. However, depression of cortical activity by anesthetics is accompanied by decreases in both glutamatergic and GABAergic synaptic transmission, with stronger effects on glutamate release^40,41^.

To directly monitor cortical GABA in a time-and depth-resolved manner, we injected AAV2/1. *hSynapsin1*.iGABASnFR into mouse primary visual cortex (V1) and measured fluorescence (with 1030 nm excitation) in a 150 μm x 150 μm x 450 μm deep volume over the course of 40 minutes. After 8 minutes, isoflurane anesthesia was administered, which is expected to suppress inhibition^41^. Fluorescence decreased 20% during anesthesia treatment and slowly returned to baseline during recovery (**Supp. Fig. S18**).

### In vivo *mouse model of epilepsy*

Having established that iGABASnFR robustly detects large-scale release events, we then tested it in small-scale, high-resolution, 2-photon neuropil imaging experiments. Focal neocortical epilepsy is typically accompanied by abnormal background electrocorticogram (ECoG) activity, including frequent inter-ictal spikes (occurring between seizures), which are dominated by GABAergic activity^42^. How seizures intermittently arise from this background is poorly understood. Numerous mechanisms have been proposed, although the strongest evidence supports a failure of feed-forward inhibition^43–46^. This could, in principle, arise because interneurons enter a state of depolarization block in the face of over-excitation and fail to release GABA^47,48^, or because shifts in the chloride reversal potential in principal cells resulting from intense GABA release render GABA_A_ receptor-mediated inhibition ineffective^49–52^. These hypotheses are not mutually exclusive, but important insights could be provided by imaging extracellular GABA during inter-ictal spiking and seizures. If extracellular GABA transients collapse rapidly at the onset of seizures, this would argue for depolarization block of interneurons, or another mechanism by which interneurons fail to be recruited or release GABA.

AAV2/1.*hSynapsin1*.iGABASnFR (or one of the two other variants) was injected into layer 2/3 of mouse V1 (**Fig. 3a-d**). Intracortical pilocarpine injection elicited focal epileptiform activity lasting up to 60 minutes^53^. This consisted of periods of inter-ictal spiking alternating with polyspikes and ECoG seizures lasting up to 1–2 minutes. During inter-ictal spiking, mice typically did not manifest abnormal motor behavior, but focal seizures were sometimes accompanied by pupil dilation and increased locomotion^54^. iGABASnFR fluorescence reliably showed transients time-locked to inter-ictal spikes (**Fig. 3e-g**). Comparing across the different sensors, iGABASnFR and iGABASnFR.F102G gave the largest fluorescence responses, while iGABASnFR.F102Y.Y137L gave lower ΔF/F. The non-GABA-binding iGABASnFR.R205A, a negative control (**Supp. Fig. S12**), was unresponsive (**Fig. 3h**), indicating that the observed fluorescence traces indeed correspond to GABA transients.

**Fig. 3.**
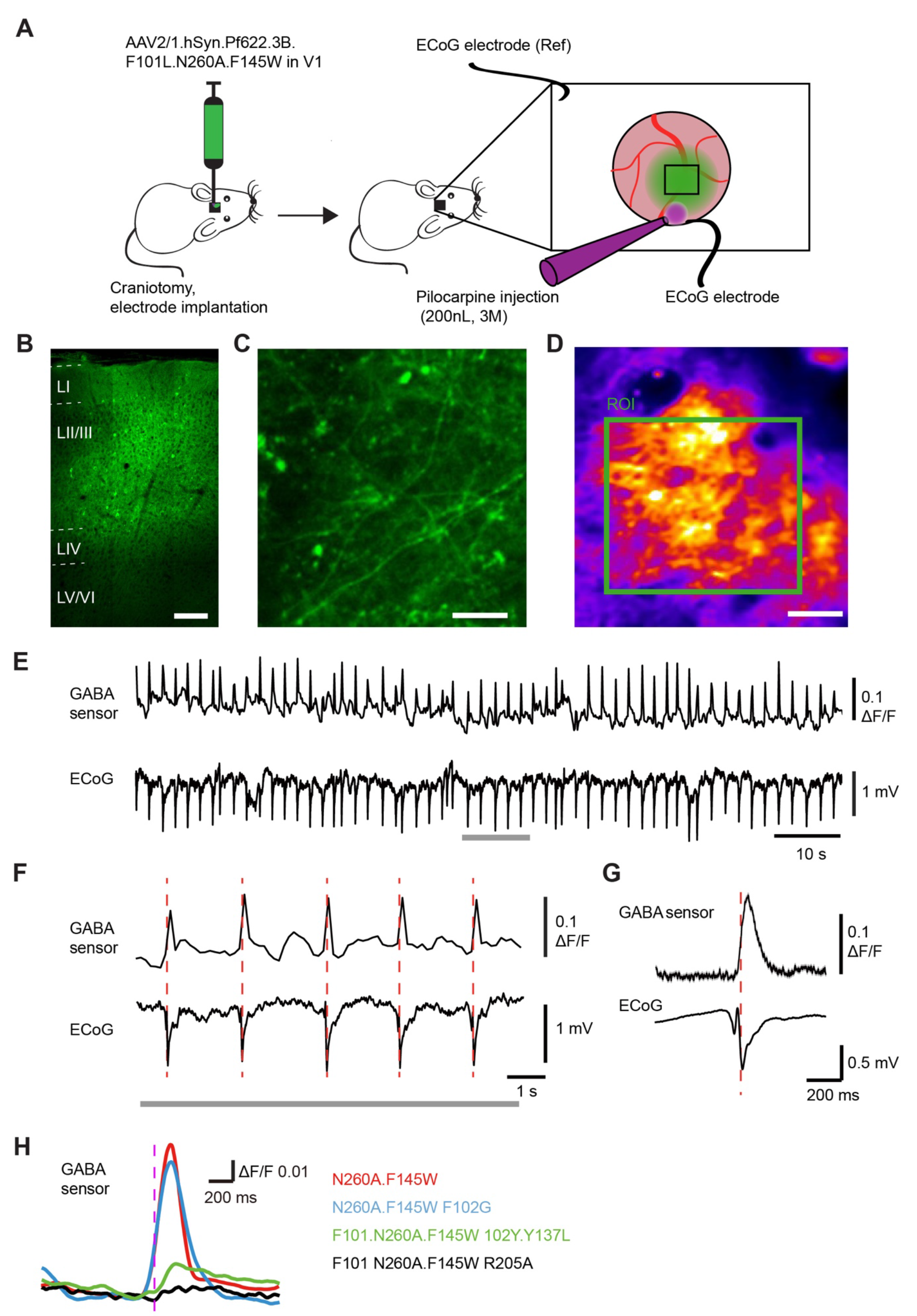
GABASnFR response during inter-ictal spiking. a) Experimental design. b) iGABASnFR expression in transduced area imaged after the end of the experiment, using immunofluorescence microscopy. Scale bar: 100 μm. c) iGABASnFR expression *in vivo* in cortical pyramidal neuron processes 100 μm below the pia. Scale bar: 20 μm. d) Average iGABASnFR fluorescence image during inter-ictal activity. Scale bar: 50 μm. e,f) Simultaneous fluorescence (averaged in green square in d) and inter-ictal ECoG. Grey bar indicates the enlargement window in f. g). Time-course of iGABASnFR fluorescence and inter-ictal ECoG averaged from 68 events in one experiment. h) Comparison of average fluorescence transients obtained with different sensors.

During periods where stereotyped polyspikes were observed, fluorescence peaks separated by <1 sec could be resolved with iGABASnFR.F102G (**Fig. 4a,b**). Discrete peaks coinciding with low-frequency ECoG components could still be seen during focal seizures lasting up to 1 minute (**Fig. 4c**). We observed a gradual decrease in fluorescence during seizures, with a slow recovery during intervals of ECoG silence in between seizures (**Fig. 4c**). This is consistent with the pH sensitivity of the GABA sensors (**Supp. Fig. S19**): a decrease in extracellular pH during seizures has been abundantly documented, and possibly factors into seizure termination^55^. The gradual increase in fluorescence between seizures is consistent with a return to normal pH over several seconds. Importantly, the slow acidification is easily separable from the GABA signals.

**Fig. 4.**
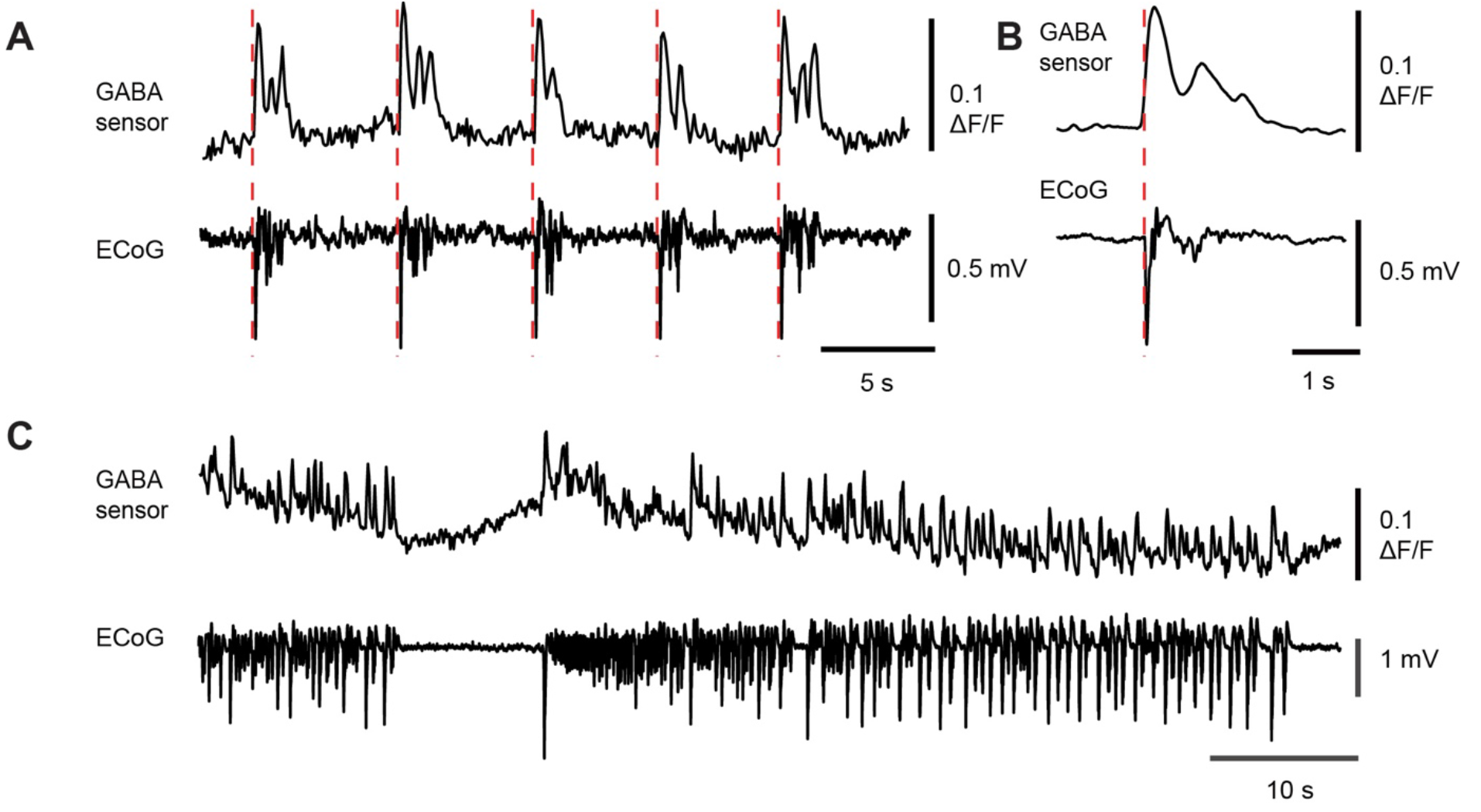
GABA sensor behavior during polyspikes and seizures. a) Trains of polyspikes (ECoG, bottom) and simultaneous iGABASnFR.F102G fluorescence (top) evoked by pilocarpine injection. b) Average time-course of sensor fluorescence and ECoG during polyspike. c) iGABASnFR.F102G fluorescence during seizures. Note that the envelope of the fluorescence transients gradually decreases during seizures, and then slowly recovers during periods of electrographic silence before the return of seizure activity.

### GABA in zebrafish cerebellum

As a final demonstration of the *in vivo* utility of iGABASnFR, we observed GABA transients in *Danio rerio* (zebrafish) cerebellum using light-sheet imaging in a fictive model of swimming^56,57^. In zebrafish, fictive swimming activity triggers robust activation of GABAergic Purkinje cells in the cerebellum^58^. We generated a transgenic zebrafish expressing the iGABASnFR.F102Y.Y137L variant under the *elavl3*/*HuC* pan-neuronal promoter (**Fig. 5a**). While the larval fish is paralyzed, and trying to swim in response to forward optic flow, its motor output to the tail muscles is captured by implanted electrodes, so that forward swimming movements can be recorded simultaneously with fluorescence imaging, with minimal motion artifacts (**Fig. 5b**). Changes in fluorescence within a cerebellar neuropil region onto which GABAergic Purkinje cells project^59^ is clear to the eye (**Supp. Movie 1**). Changes in fluorescence peaked roughly 200 msec after the onset of swimming (**Fig. 5c**), with this region of the cerebellum producing upwards of 8% changes in fluorescence. This result indicates that iGABASnFR can reliably detect GABA transmission that are potentially critical for the motor control of zebrafish.

**Fig. 5.**
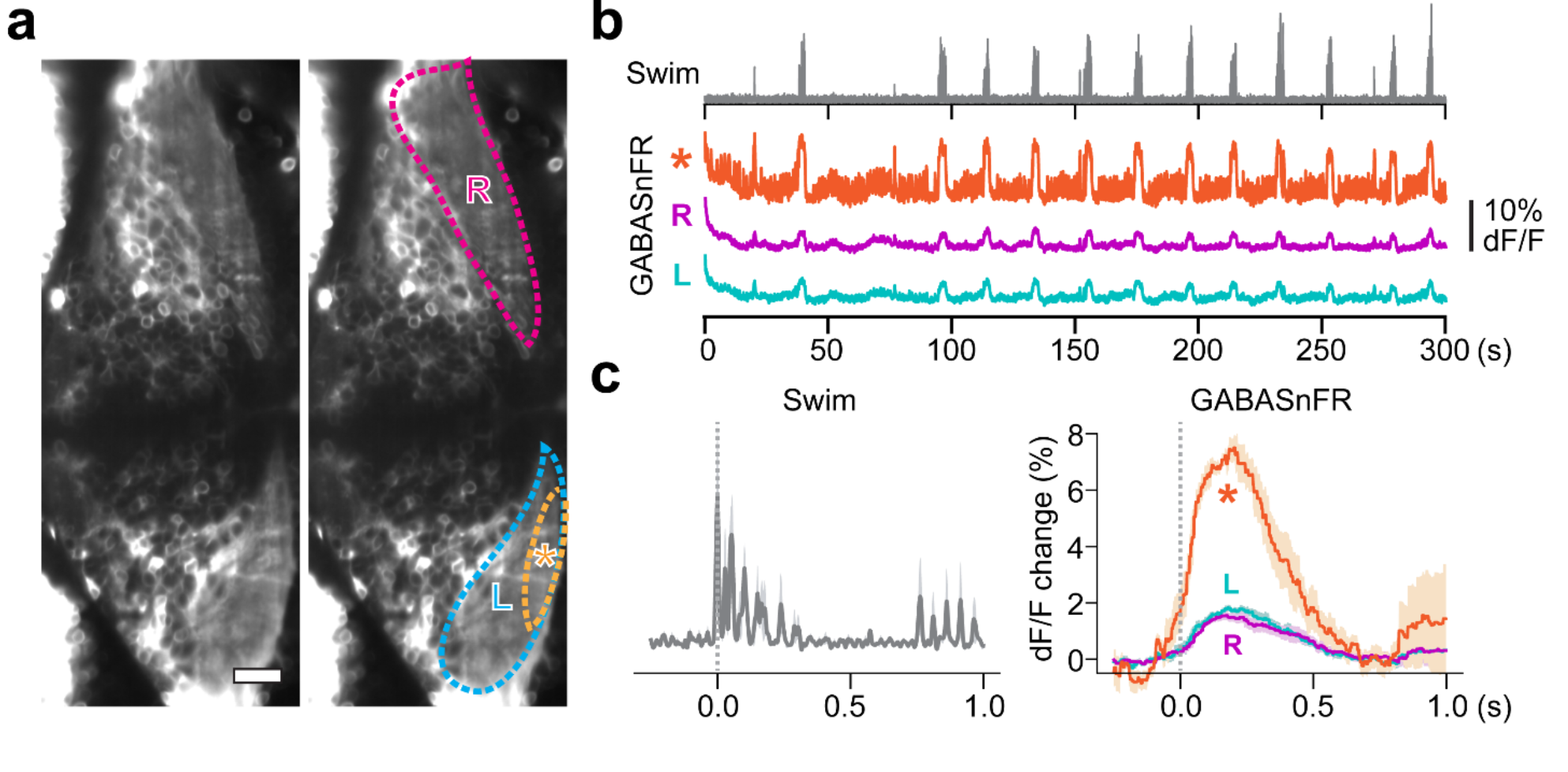
GABA response seen with iGABASnFR.F102Y.Y137L fluorescence in a fictive model of swimming in zebrafish larvae. See also Supp. Movie 1, in which changes in fluorescence within the region denoted by an asterisk* are obvious. a) Light sheet image of zebrafish cerebellum expressing iGABASnFR.F102Y.Y137L under the *elavl3/HuC* promoter. b) Motor signals (grey) and fluorescent signals from ROIs indicated in (a). c) Motor signals (left) and fluorescence changes (right) averaged across 5 swim events. Shadows represent s.e.m. across swim events.

## Discussion

To understand neural circuit function, numerous specific inputs onto neurons must be disentangled from their integrated output. Current sensor technologies, namely iGluSnFR^12,25^, GCaMP6^11^ and jGCaMP7 (unpublished), and jRGECO^10^ are sensitive enough to allow reliable detection of excitatory synaptic transmission, excitatory post-synaptic currents, and action potentials. Indicators for inhibitory synaptic transmission and inhibitory post-synaptic currents (IPSCs) have lagged far behind. iGABASnFR offers the best performance of existing GABA indicators, and is the only one appropriate for *in vivo* use. Calcium and glutamate sensors have been iteratively optimized, both to improve overall performance and to match to precise requirements of specific settings. We expect further development of iGABASnFR, with altered affinity, kinetics, and improved signal-to-noise ratio. Different colors of iGABASnFR could also be developed, allowing simultaneous imaging in orthogonal chromatic channels, as has been done with iGluSnFR^25,60^.

We hope that the demonstrated use of iGABASnFR in models of epilepsy and in the cerebellum of fictively behaving zebrafish will facilitate a better understanding of the role of GABA in various circuits, during development^61^, and in disease states. Aberrations in GABAergic signaling have been demonstrated in Alzheimer’s disease^62^, Parkinson’s^63^, Huntington’s^64^, schizophrenia^65^, and autism/ Rett syndrome^66^. iGABASnFR imaging in appropriate animal models will facilitate greater understanding of these mechanisms. iGABASnFR could also allow easy screening^67^ of candidate transporters such as the vertebrate mitochondrial GABA transporter^27^. Applications outside of neuroscience are also possible, such as separating the critical roles of GABA in plant metabolism and signaling^68^.

## Acknowledgements

We would like to thank Catherine S. Nicholson-Guthrie (Indiana University) for the gift of the *Pseudomonas fluorescens* strain CNG89. David Stern, Andy Lemire and Damian Kao for helping sequence the genome of CNG89. Deepika Walpita for rat neuronal culture. And John Macklin & Ronak Patel for collecting 2-photon spectra.

YS, ML and DMK are supported by the Medical Research Council and Wellcome Trust. VM is supported by Epilepsy Research UK.

